# Solar Induced Fluorescence Retrieved from Avocado (*Persea americana* Mill.) Canopies Along the Season Correlates with Sugar Levels in the Developing Fruit

**DOI:** 10.1101/2023.07.24.550262

**Authors:** S. Vitrack-Tamam, H. Yasour, D. Hamus-Cohen, R. Erel, L. Rubinovich, O. Liran

## Abstract

Solar Induced Fluorescence (SIF) emitted from the photosynthetic apparatus is related indirectly to biomass production. It can be remotely sensed from airborne platforms, yet, the spatial resolution from satellites, for example, is too low for commercial agricultural uses. We used a non-imager point spectroradiometer mounted on SIF-UAV (Unmanned Aerial Vehicle) system, and retrieved SIF at high spatial resolution, which is fitted for precision agriculture applications. In this study, we tracked the spatial variation of SIF along the season over a commercial avocado (P. americana Mill) orchard. The retrieved SIF signals were Krieg-interpolated to create a continuous SIF layer that allowed the estimation of SIF values for each tree in the orchard. We show that the SIF signal retrieved over the canopies in the main fruit development stage is highly correlated with the levels of sugars accumulated in the ripening avocado fruit. It established the foundations for high-spatial resolution detection of the natural variation of photosynthesis activity, which may lead to in-season adjustments of agronomic inputs using precision agriculture technologies.

## Introduction

Various climate models predict that the average global temperature will continue to increase between 1.4 °C and 5.5 °C into the next century (Masson-Delmotte et al., IPCC2018). Global warming leads to climate change which results in attenuated geographical precipitation coverage, prolonged heat waves and frequent drought events that are detrimental to agricultural production. Together with the expected increase in world population, climate change demands the development of improved agricultural practices and management tools at various scales to secure optimum crop yield. Passive optical remote sensing has been used for many years for deducing crop status and canopy parameters. It includes measurements from ground, aircraft and satellite (Maier, Günther, and Stellmes 2004). Vegetation Indices (VIs) such as Normalized Differential Vegetation Index (NDVI) are sensitive to canopy chlorophyll levels and as a result represent a quantity of the absorbed photosynthetically active radiation (Gitelson, Merzlyak, and Lichtenthaler 1996). While this and other reflectance-based vegetation indices might be related to the product of photosynthesis activity, they are unable to detect the response of photosynthetic physiology to temporal attenuations in environmental parameters. A search for another, much more sensitive index resulted in Solar Induced Fluorescence (SIF) as an index of photosynthetic activity and biomass production (Porcar-Castell et al. 2014). Not all the light which is absorbed by the Light Harvesting Complexes (LHC) of a plant is used to synthesize sugars (Baker 2008). Part of it is dissipated as heat, and the rest is emitted back to the environment as a fluorescence. The emitted fluorescence is linked to photosynthesis (Kitajima and Butler 1975). SIF was found to report not only on primary productivity, i.e. gas exchange and carbon assimilation on a global scale (Mohammed et al. 2019), but also on the performance of the photosynthetic system of crops grown within an agriculture field (Zarco, 2016).

There are several remote sensing retrieval techniques to extract SIF from a crop reflectance signature, to name a few: Extraction of the fluorescence addition to the reflected light at 762 nm, where oxygen absorbs sunlight (Maier, Günther, and Stellmes 2004), and at the red-edge range (Pablo J. Zarco-Tejada et al. 2003). With improvements in sensor capabilities, it is now available to retrieve SIF from upwelling radiance returning from crops with ultra-sensitive point spectroradiometers (Burkart et al. 2013). The technique makes use of recording the oxygen absorption line at 759.39 nm in a very high spectral resolution (<1 nm Full Width at Half Maximum, FWHM). Then, when compared to a white reference, by using the Fraunhoffer Light Discriminator technique (FLD, (Plascyk 1975)), fluorescence signals of crops performing photosynthesis were shown to relate to crop production (Rascher et al. 2009), temporal dynamics of photosynthesis gas exchange signals (Cogliati et al. 2015), and water stress (Daumard et al. 2010). Recently, non-imaging spectroradiometers were mounted on an Unmanned Aerial Vehicles (UAV) in order to track photosynthetic activity over agriculture fields on high spatial and temporal resolutions. Comparison between SIF imager and SIF retrieved from a non-imaging spectroradiometer shows high correlation between SIF retrieved in both methods (Wang et al. 2022). SIF retrieved from a UAV was also correlated with leaf chlorophyll a fluorescence parameters and water stress in potato crop in an open field (Xu et al. 2021).

Avocado (*Persea Americana* Mill) is a commercially important fruit tree due to its highly beneficial nutritional values (Rendón-Anaya et al. 2019). Avocado is a climacteric fruit, which means that it increases its respiration and ethylene production during ripening, and can continue ripening after harvesting, which results in a fruit with a low shelf life (Chen et al. 2017). For this reason, a pre-harvest understanding of parameters such as photosynthetic activity and biomass production are important to assess harvest time (Robson, Rahman, and Muir 2017). As sugar content in the fruit is attenuated along the ripening period (Delisiosa 1992) it may be related to photosynthetic cues that can be measured with remote sensing techniques. Implementing a remote sensing technique which is able to track photosynthetic activity directly from the canopies of the trees and then report on biomass production on the fruit level will shed a light on the intricate mechanism of sugar transfer from sources to sinks. The objective of this study was to relate canopy extracted SIF signals measured using a non-imaging spectroradiometer mounted on a UAV with sugar content in fruits of Avocado cv. ‘Hass’. Monthly SIF scans using a UAV and fruit sampling were performed during the main fruit growth stage.

## Materials and Methods

### Experimental orchard

A commercial avocado (*P. americana* Mill.) cv. ‘Hass’ orchard was selected for this study. The orchard is 11 years old and located near Kfar Glickson) 32°29’55.9"N 35°00’08.4"E (, Israel. The selected plot in the orchard consists of 31 rows with 43 trees per row. Tree spacing is 4 m between rows and 4 m between trees.

### Field campaign

High resolution reflected spectra were acquired using a drone, once a month, from July to November (Table 1), which is the main fruit growth stage. In each flight date, representative fruits were picked from trees of three different crop loads (0-10, 12-40, 60-80 kg tree^-1^, for empty, half loaded and loaded trees, respectively). For each sampling date, one fruit from each of six different trees of each of the crop load groups was randomly selected and used for the determination of the content of various sugars. In total, 18 fruits were selected for each of the sampling dates.

**Table 1.** Meteorological conditions in each of the spectral acquisition sampling dates along the season.

| Date<br>(dd,mm,yy) | Flight time<br>(hr) | Irradiance<br>(W m<sup>-2</sup>) | Relative<br>humidity<br>(%) | Air<br>temperature<br>at 2m (C<sup>0</sup>) | Wind Speed<br>(m s<sup>-1</sup>) |
| --- | --- | --- | --- | --- | --- |
| 04.07.21 | 10:00 | 566 | 61 | 29.7 | 5.1 |
| 04.08.21 | 10:00 | 770 | 58 | 33.3 | 2.2 |
| 12.09.21 | 10:30 | 642 | 58 | 28.3 | 3.2 |
| 03.10.21 | 12:10 | 656 | 59 | 28 | 3.8 |
| 14.11.21 | 10:00 | 599 | 18 | 25.7 | 6.6 |

### UAV system platforms

Mosaiced RGB pictures, providing an aerial view of the orchard, were created with MAVIC2-Pro (DJI, Shenzen, China) using its default 20 Mega Pixel camera. The drone flew at a height of 50 m above ground, at a speed of 5.5 m s^-1^. Images were loaded into Pix4D-React (Pix4D, Prilly, Switzerland) in order to create the continuous image layer. The image layer was used only for presentation of the data. In order to construct continuous layers of spectral indices, MATRICE 600-M drone (DJI, Shenzen, China) equipped with a GPS-guided gimbal Ronin-MX (DJI, Shenzen, China) was used. Point spectroradiometer, a card computer and a battery (STS-VIS developer kit, Ocean Insight, FL, USA) were mounted onto the gimbal. The point spectroradiometer was always faced down at nadir, except during take-off and landing, when it was folded perpendicular to the nadir. The drone flew at a height of 15 m above the canopies of the trees (∼20 m above ground) at a speed of 3.7 m s^-1^. This resulted in covering an area of approximately 6.5 m in diameter per each spectral acquisition.

### Spectral acquisition

#### Spectral acquisition system

Spectral acquisition was performed as described in Liran (LIRAN 2022). Two STS-VIS micro-spectroradiometers (STS-VIS Developer kit, OceanInsight, Largo, FL, USA) acquired the spectra simultaneously - one was mounted on the drone which retrieved upwelling radiance from the crops (AIR unit) and the second was mounted on a stand at the ground retrieving upwelling radiance from a reference plate (94% reflectance plate, Permaflect^®^, LabSphere, USA) (GROUND unit). Each spectroradiometer was radiometrically calibrated according to the manufacturer instructions with a NIST (National Institute of STandards) calibrated halogen lamp (HL-2000-LL, OceanInsight, FL, USA).

#### Spectral cross validation between the two sensors units

Cross validation of the two units measuring radiometric signatures was performed on the Fraunhofer O_2_-A absorption band acquired from the 94% white plate used as a reference. Essentially, both units’ spectra minimum of the O_2_-A line was set at 759.37 nm. By placing the two spectra of the AIR and GROUND units one on top the other, O_2_-A absorption line was checked to obtain the same characteristics – height, width and size.

#### Spectra acquisition and calculations

Spectra were transferred from machine units (photon counts) to flux units via the following formulation according to the manufacturer’s suggestions:

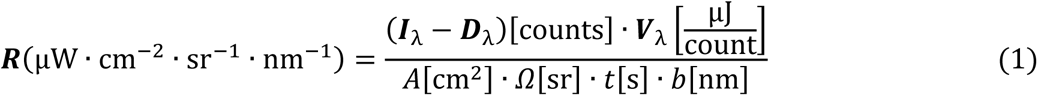

Where, ***R*** is the light flux radiance incoming to or upwelling from the target, ***I***_λ_ is the photon counts in each wavelength, ***D***_λ_ is the dark current values in each wavelength (Burkart et al. 2013), ***V***_λ_ is the calibration vector of the spectroradiometer, *A* is the area from which photons are returning into the pupil of the spectroradiometer, *Ω* is the solid angle of the cone from which photons are collected, *t* is the integration time selected during a single acquisition, and *b* is the bandpass, the spectral width between two successive wavelengths in the recorded spectrum. The discrete difference value between successive wavelengths varies from 0.4 to 0.45 and should be therefore treated as:

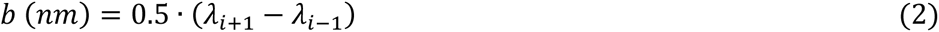

Normalized reflectance spectra from each pair of reflected energy flux spectra acquired simultaneously by the AIR and GROUND units was created according to Gordon and Wang (1994) (Gordon and Wang 1994):

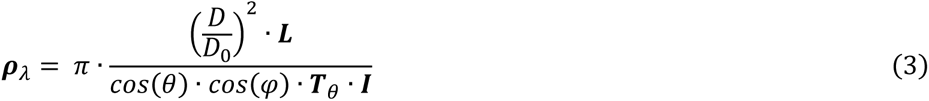

Where, **ρ**_λ_ is the normalized reflectance vector comprised of a ratio of the upwelling radiance flux (***L***) and incoming irradiance flux (***I***), multiplied by constants which relate to the distance of the earth from the sun (*D, D_0_* – distance at the day of the measurement and the average distance, respectively), the position of the sun in the sky (*θ,φ* are the sun elevation and azimuth, respectively), and ***T****θ* is the atmospheric diffuse transmittance for each wavelength and the position of the sun in the sky. This was estimated separately using simulation of the irradiance during the acquisition date and conditions, with MODTRAN software (Spectral Sciences, Burlington, MA, USA) (Berk, Bernstein, and Robertson 1987).

### Flight log transformation

Each flight log file was downloaded directly from the Matrice 600 M drone and converted to a data file (*.csv) with the phantomhelp.com website (https://www.phantomhelp.com/logviewer/upload/). Then, the geodesic ESPG::4326 coordinate system format was converted to a 2D projected coordinates on the mosaic image in format of ESPG::32636 via OSgeo package (GDAL/OGR contributors 2022) written in Python language (Rossum, Fred, and Drake Jr 1995).

### Alignment of flight and sensor data

The exact position of each spectral acquisition was extracted from the flight data and included the GPS transponder point and its time of recording. On the other hand, the time of acquisition by the spectral sensor was extracted from the developer kit. The clocks of the drone and the sensor are not synchronized and require an alignment procedure. A prominent, single GPS data point, corresponded to an "easy to identify" spectral measurement point and was designated as an “anchor point”. It was found that a spectral reflectance of soil is an "easy" target to be decided as the "anchor point" - the first geographical point just before Normalized Differential Vegetation Index (NDVI) (Rouse Jr et al. 1974) reaches a value above 0.8, i.e. the transfer from soil GPS point to a canopy point. The drone logged the GPS data at different starting and interval time points than the sensor. In order to synchronize the two clocks, the following alignment procedures were executed:

a. Coarse alignment – only the actual flight segment above the trees was considered.
b. Fine alignment – algorithm presented in (Figure 1) was used in order to align the spectral measurements with the respective GPS points.

**Figure 1.**
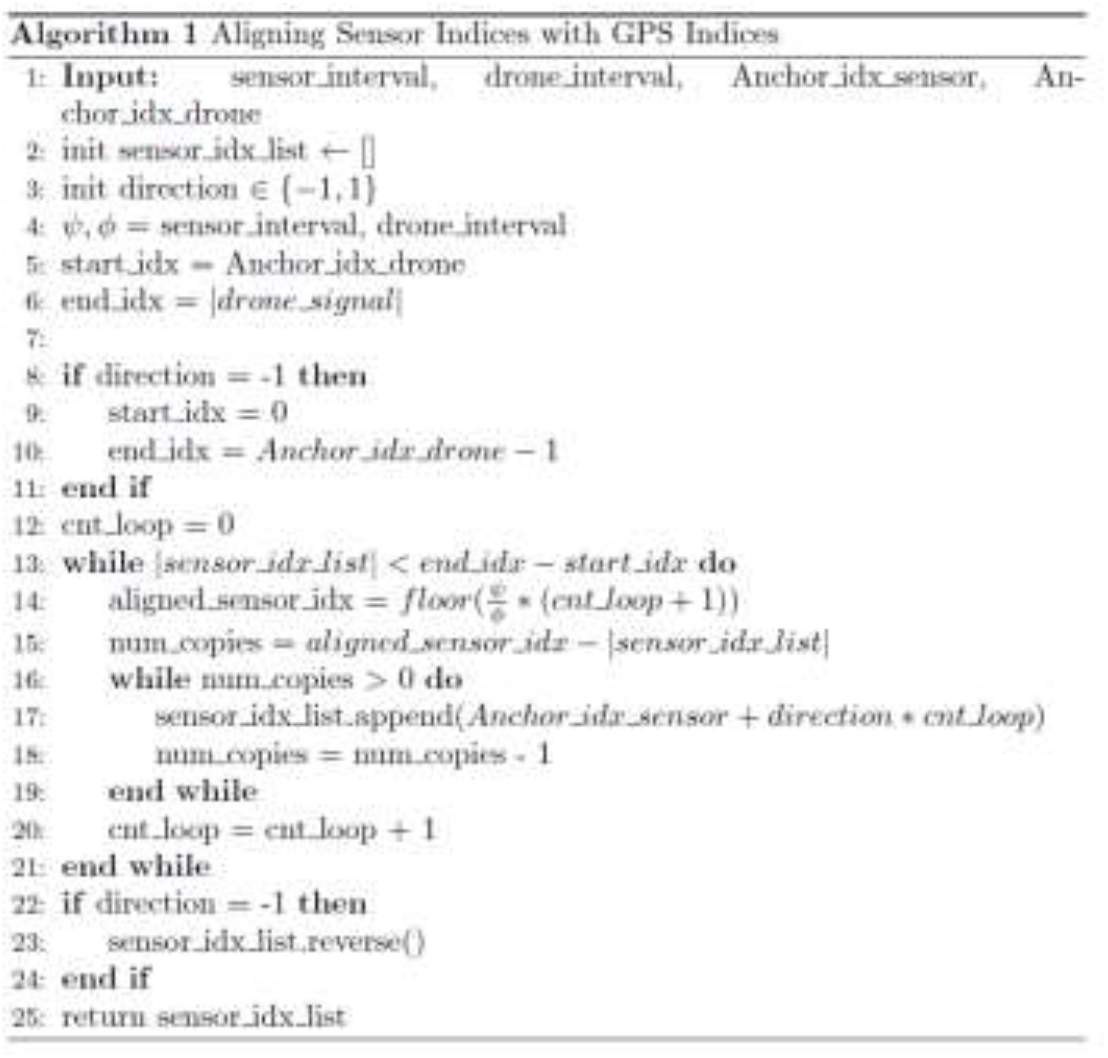
Code segment of aligning drone GPS and Sensor’s indices together.

This step was verified manually: at the end of each drone pass, the turn command was performed above a soil with NDVI value <0.3. Therefore, these low NDVI points found at the end of each pass and the borders of the orchard were used as the manual cross validation that the map spectral points were aligned correctly with the actual geographical area.

### Vegetation Indices (VIs) calculation

Two canopy structure VIs and two SIF indices were calculated from either the reflectance spectra, or from the radiant energy fluxes from the canopy, respectively (Table 2).

**Table 2.**
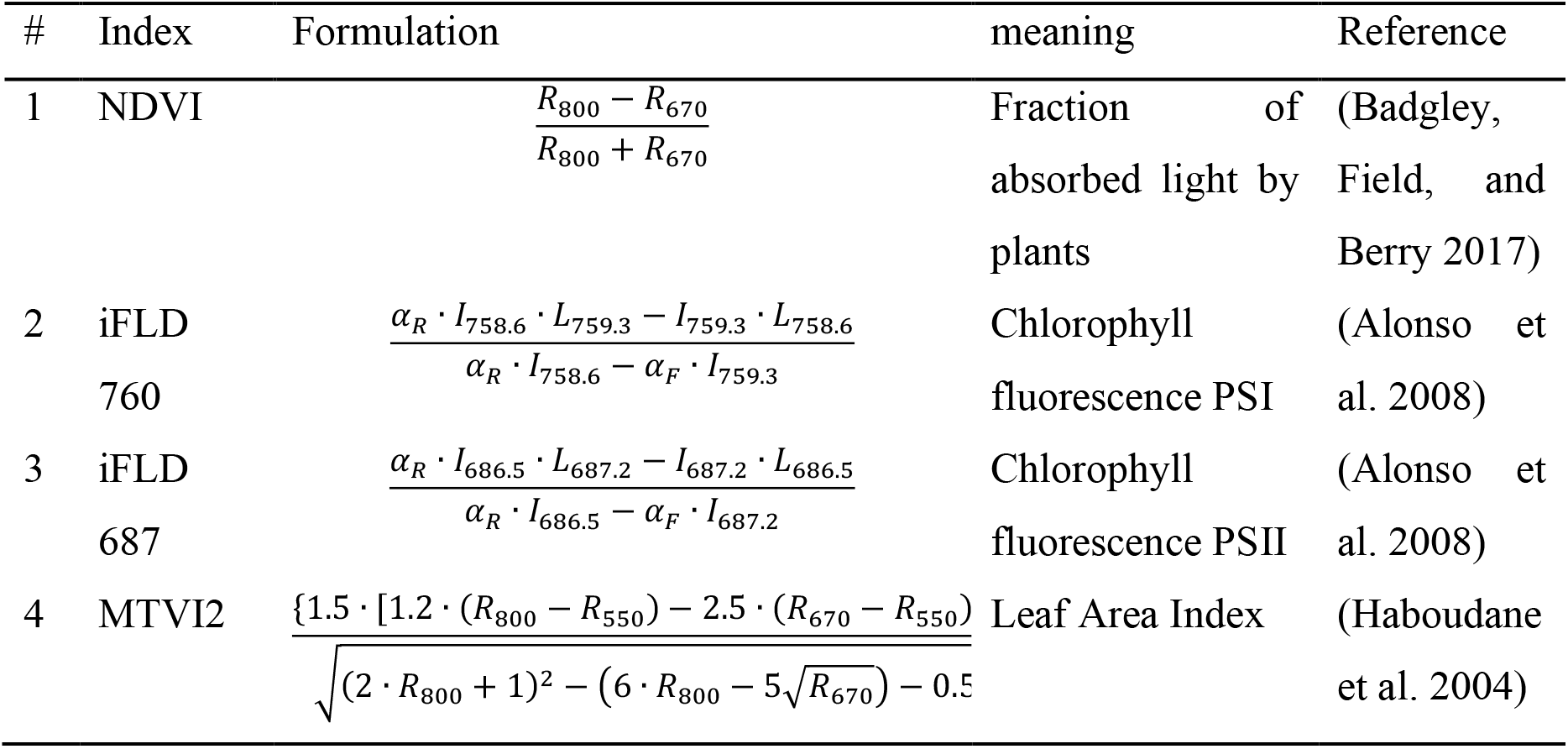
List of vegetation indices formulations used in this study.

### Data allocation per tree

Data from each spectral acquisition was interpolated using the Kriging technique with Variogram Estimation Toolbox in Python language (Mälicke 2022). This is because the drone movement above the field in each pass was not always aligned with the tree lines, and not always accurate due to winds. Therefore, an interpolated continuous layer was constructed. The semivariance estimator used was of Matheron et al. (Matheron 1963) with an exponential fit (Journel and Huijbregts 1976). The resulting continuous predicted layer was cast on top of the experimental area RGB picture according to its geotagged location. In order to assess each tree location within the designated area, each tree canopy was referred to as a perfect circle with a diameter of 4 meters. These circles were aligned with the actual position of the trees. However, within the orchard, some of the trees are not situated exactly on spot, and therefore in order to extract the values for each tree on the map as accurately as possible, the actual place was approximated according to an even spacing along each row by considering the total number of trees in that row:

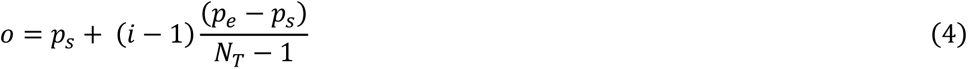

Where, *o* is the position of an arbitrary tree, *p_s_* and *p_e_* are the positions of trees at the start and end of each row, *i* index of the tree and *N_T_* is the number of all the trees in each row. Finally, the value of indices for each tree was calculated as the average of all the values that fall within the 4 m diameter of the circle of the canopy.

### Sugar contents of avocado fruits

Fruit sampling for metabolomics analysis was conducted on the same day of spectral acquisitions analysis (Table 1). In this study the target was to relate between vegetation and photosynthetic indices retrieved from the canopies (leaves) to hexose and heptulose sugars in the fruit’s tissues. Frozen avocado fruit pericarp was ground to a fine powder in liquid nitrogen using mortar and pestle. Sample preparation and metabolite extraction were performed according to a modified method described by Roessner-Tunali et al. (Roessner-Tunali et al. 2003): 100 mg of frozen pericarp powder was ground with a bead beater (MillMix 20, Domel, Železniki, Slovenia) for 45 s at 24 Hz with 700 μl of cold methanol and 40 μl of internal standard (ribitol (A5502, Sigma-Aldrich, St. Luis, MO, USA), 0.2 mg in 1 ml of water). The samples were vigorously shaken for 20 min at 4°C and then 750 µL water was added, and the mixture was vortexed and centrifuged at 13,000g at 4°C. 400-µL aliquots were taken and dried out using Christ Alpha RVC (Gemini BV, the Netherlands). The dried-out material was re-dissolved and derivatized for 90 min at 37°C (in 40 µL of 20 mg mL-1 methoxyamine hydrochloride (89803, Sigma-Aldrich, St. Luis, MO, USA) in pyridine (270970, Sigma-Aldrich, St. Luis, MO, USA) followed by a 30-min treatment with 70 µL MSTFA [N-methyl-N(trimethylsilyl)-trifluoroacetamide (69479, Sigma-Aldrich, St. Luis, MO, USA) at 37°C and centrifugation. Following derivatization, samples were analysed in an Agilent 6850 GC/5795C mass spectrometer (Agilent Technology, Palo Alto, CA, USA). Sample volume of 1 μl was injected as a 1:50 split into the GC column. Helium was used as carrier gas at a flow rate of 1 mL min-1. Chromatography was performed using an HP-5MS capillary column (30 m × 0.250 mm × 0.25 μ m) (Agilent J&W GC Columns, 9091S-433E, Santa Clara, CA, USA). Mass spectra were collected at 2.4 scan s-1 with an m/z 50–550 scan. The temperature program was according the following: hold at 50°C for 5 min, followed by ramping up the temperature to 270°C with ramp rate of 7°C min-1, and followed by a cleaning stage (increasing the temperature to 320°C with a ramp rate of 10°C min-1; hold at 320°C for 10 min). Both ion chromatograms and mass spectra were evaluated using MassHunter software (Agilent, Santa Clara, CA, USA). Sugars and were identified by comparison of retention time and mass spectra with authentic standards (Sigma, Sigma-Aldrich, St. Luis, MO, USA), while other metabolites were identified using NIST14 (Agilent, Santa Clara, CA, USA).

### Statistical analysis

We followed the statistical analysis that was presented in (Liran et al. 2020). Each group in the study (fruit load by date) was checked for the presence of outliers where values found 1.5 times outside the interquartile range were omitted from further analysis. Then, the groups were checked for normal distribution with Shapiro–Wilk’s test and for homogeneity of variances via Levene’s test. Analysis of variance (ANOVA) was applied when both tests were satisfied. In case the homogeneity of variances test was violated, Welch’s ANOVA was used instead, and if Shapiro-Wilk’s test was violated, a Kruskal– Wallis ANOVA was used. In order to compare between different dates, a repeated measures ANOVA was used if Mauchly’s sphericity test was satisfied. In case the sphericity test was violated, Friedman’s non-parametric repeated measures ANOVA was used instead. Statistical significance was always set at p < 0.05. Correlation matrix was constructed with Spearman’s non-parametric formulation because the values of each of the metabolites’ content was attenuated at least three orders of magnitude along the season (Myers and Sirois 2004). Autocorrelation between sampled groups for the vegetation indices were checked using the Moran’s autocorrelation coefficient statistically significance index (Moran 1950). Then, the coefficients were transformed to Z-score and checked for statistical significance. In case they were found true, those groups were taken out of the correlation analysis, but were marked as autocorrelated in analysis afterwards (Afyouni, Smith, and Nichols 2019). Data analysis was performed in R studio (RStudio Team 2022), in R language (R Core Team 2021).

## Results

### Construction of vegetation indices’ values for each tree in the orchard

Sensor’s position in the field was approximated by calculation (Figure (2a)), where the anchor point stitching together the information from the drone’s GPS and the sensor acquisition in place and time was selected to be the first point in the spectrum just after passing from soil sample to the canopy sample (Figure (2b)). The reflected spectra covered various types of objects in the scene: soil only as 0-0.3 NDVI, canopy only NDVI > 0.8, and a mixture of both as 0.4 - 0.7 NDVI, thus the information was divided into three categories: soil, soil+ canopy, and canopy (Figure (2c) α, β and γ designations, respectively). Finally, the information was casted upon a mosaiced geotagged RGB picture of the area (Figure (2d)), where we validated the correctness of the synthesis of the data visually.

**Figure 2.**
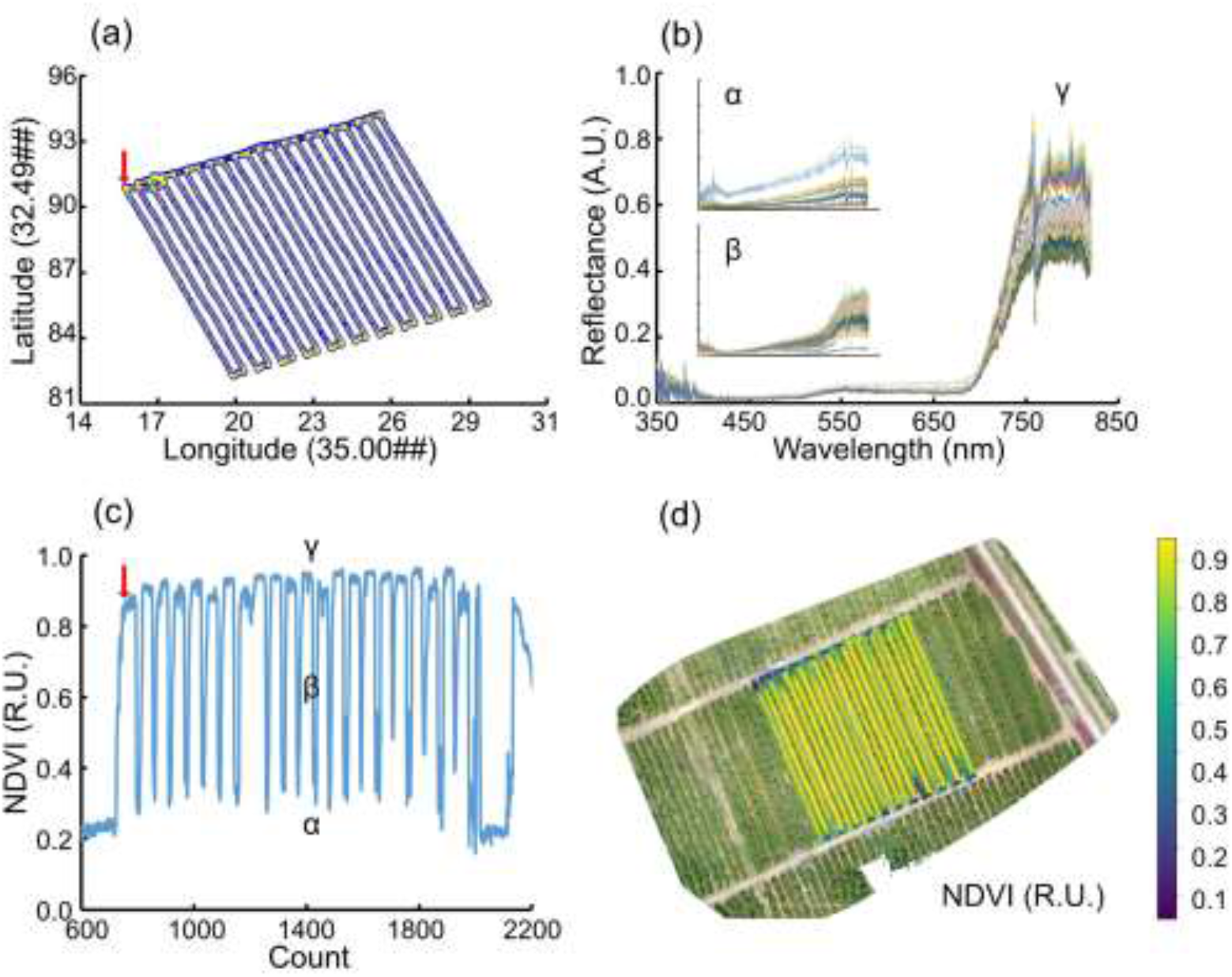
Synthesis of information from the spectroradiometers and the drone’s flight path. (a) Projection of the drone’s flight path onto a 2D plane in ESPG::32636 coordinates system (>8000 Blue big dots in the plot). (b) Reflectance spectra acquired during the flight over soil, soil + canopies of trees and canopies of trees only (α, β and γ, respectively). The subset units on the XY scales were omitted due to lack of space in the figure. (c) Calculation of NDVI vegetation index from each of the acquision points. The three spectral groups of acquired objects are marked on the plot. The red bold arrow indicates the "anchoring point" for the matching algorithm between the drone and sensor time recordings. (d) Projection of the NDVI vegetation index on the mosaiced RGB picture of the experimental area. Colors from dark purple to bright yellow are a continuous representation of values from 0.1 (soil only) to 0.9 (vegetation only) NDVI relative units.

Gaussian interpolation (Kriging procedure) was generated for each of the four indices calculated in this study in order to approximate their value for each tree in the orchard. For example, the NDVI-approximated layer for one of the dates is presented in Figure 3. Kriging resulted in a zebra strip sub-areas (Figure (3a)). These alternating strips show sub-areas with NDVI 1.0 values, and between them NDVI 0.7-0.9 mixed sub-areas. The dark colours at both ends of the transverse lines represent the NDVI values of 0.2-0.4 (mixed soil/vegetation and soil only), due to the drone crossing the border between the vegetation and the soil objects. The interpolated NDVI values for each tree in the orchard are presented in Figure (3b), thus it can be correlated with the sugar contents in fruits that were sampled from trees that their geo tagged location was known.

**Figure 3.**
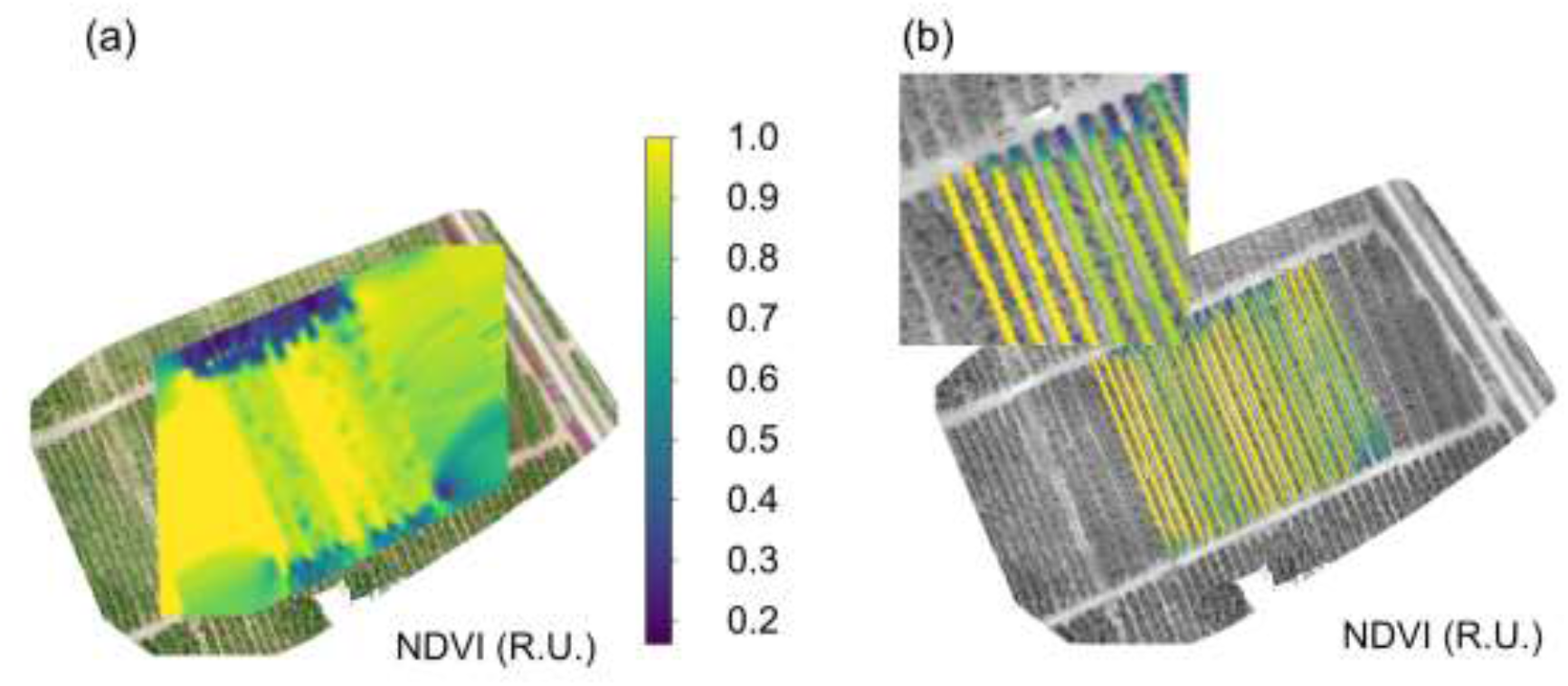
Continous layer of a vegation index casted on the geographical map. (a) Result of kriging interpolation over the whole flight area with an NDVI vegetation index. Colors of deep purple to bright yellow represent NDVI values from 0.1 to 1.0, respectively; (b) Approximated representation of trees values performed on the kriging layer.

### Seasonal dynamics of canopy vegetation indices, canopy SIF indices and sugar content

Sugar depletion in avocado fruits was documented as the season progressed after June’s drop (Figure 4). Sucrose and fructose levels were attenuated more than three orders of magnitude during the pass from July to August (Figure (4a) and (4b)- note the logarithmic Y scale). In the case of sucrose and fructose, there was only a difference between July and the rest of the season (Figure (4a), (4b)). The two C_7_ sugars D-mannoheptulose and perseitol show exponential decay trend, similar to sucrose and fructose (Figure (4c), (4d)). The levels of the hexose sugars present the same logarithmic decay along the season, albeit the magnitude of the decrease was smaller than the sucrose and fructose. There was no visual gradient of sucrose content between the three fruit loads within each date, although there was a trend of decreasing sugar content with increasing fruit load in September and October (Figure (4a)). Fructose content showed no trend with increasing fruit loads (Figure (4b)). Statistically significant increase in C_7_ sugar content with increasing fruit load was apparent only in October.

**Figure 4.**
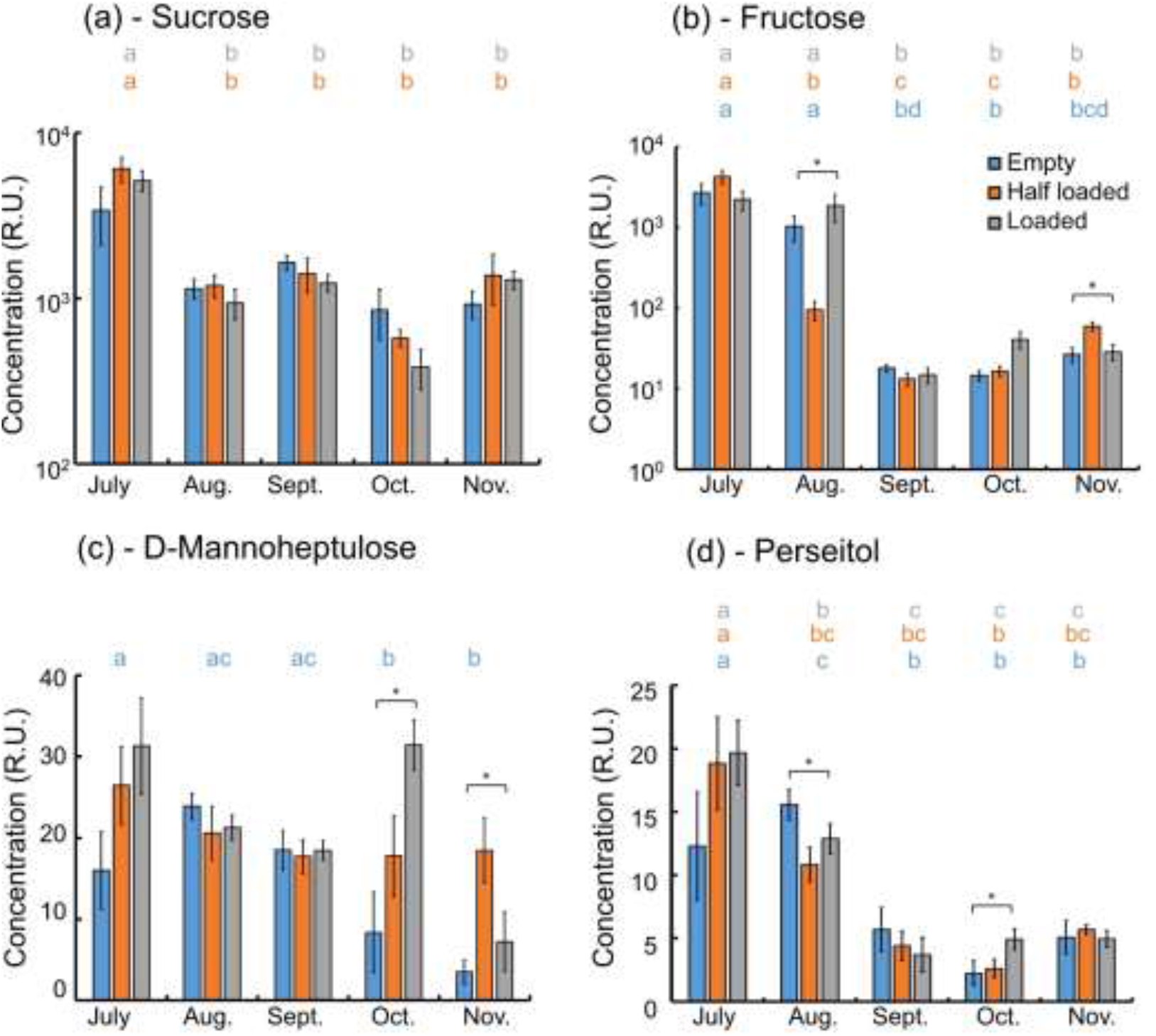
Seasonal variation in sugars levels in ripening ‘Hass’ avocado fruits. Panels (a), (b), (c), and (d) represent Sucrose, Fructose, D-Mannoheptulose and Perseitol, respectively. In sucrose and fructose, the values are transformed to natural logarithm and in perseitol and D-mannoheptulose the values are divided by 1000 for ease of representation. Fruit load groups are represented in colors, where empty (0 – 10 kg tree^-1^), half loaded (12 – 40 kg tree^-1^), and loaded (60 – 80 kg tree^-1^) are colored in blue, orange and grey, respectively. Letter annotation represents statistically significant differences between dates. Statistically significant differences were always determined at p < 0.05. N=6, error bars represent standard error of the mean (S.E.). Asterisks represent statistically significant differences.

A correlation analysis was performed between NDVI, MTVI2, SIF687 and SIF760 indices and the levels of the four sugars measured in the fruit pericarp. Spatial autocorrelation was calculated for each tree within the crop load groups as a prerequisite for independence between samples in a geographical area. This is because autocorrelation treats independent biological samples as technical repeats, i.e. inherently correlated (Legendre, 1993). Therefore, the analysis of the Moran index which determines how much autocorrelation exists between samples taken from different geographical positions was performed on the calculated indices before the kriging interpolation (Table 3). In general, the autocorrelation was negative for almost all the fruit load groups across all dates, meaning that there is a homogenous distribution of tree groups within the orchard, rather than two or three clusters of similar nature. The values of the Moran index declined over time, from ca. -0.6 on average during the summer to half of that during the winter, close to the time of the commercial harvest. Statistically significant autocorrelation was more common towards the end of the season, because towards the end of the season, most of the orchard reaches a uniform stage where most of the sugars in the fruit are consumed (Figure 4).

**Table 3.** Statistical significance of Moran’s index of spatial autocorrelation for each vegetation index calculated in this study, in each fruit load and for each month during the growing season, starting after June’s drop.

| Month | Fruit load | MTVI2 |  | NDVI |  | iFLD687 |  | iFLD760 |  |
| --- | --- | --- | --- | --- | --- | --- | --- | --- | --- |
|  |  | Moran | P-value | Moran | P-value | Moran | P-value | Moran | P-value |
| July | Empty | -0.43 | 0.02* | -0.54 | 0.00* | -0.52 | 0.00* | -0.54 | 0.00* |
|  | Half loaded | -0.10 | 0.14 | -0.31 | 0.07 | -0.37 | 0.01* | -0.39 | 0.01* |
|  | Loaded | -0.71 | -- | -0.74 | 0.50 | -0.62 | 0.50 | -0.60 | 0.50 |
| August | Empty | -0.17 | 0.11 | -0.58 | 0.03* | -0.56 | 0.05 | -0.54 | 0.06 |
|  | Half loaded | -0.07 | 0.08 | -0.31 | 0.10 | -0.42 | 0.02* | -0.42 | 0.02* |
|  | Loaded | -0.23 | 0.36 | -0.35 | 0.06 | -0.28 | 0.20 | -0.51 | 0.00* |
| September | Empty | -0.41 | *0.01 | -0.24 | 0.31 | -0.26 | 0.26 | -0.37 | 0.02* |
|  | Half loaded | -0.18 | 0.26 | -0.62 | 0.01* | -0.61 | 0.01* | -0.57 | 0.02* |
|  | Loaded | -0.59 | 0.50 | -0.60 | 0.50 | -0.59 | -- | -0.61 | 0.50 |
| October | Empty | -0.52 | 0.02* | -0.57 | 0.01* | -0.36 | 0.23 | -0.52 | 0.02* |
|  | Half loaded | -0.32 | 0.26 | -0.45 | 0.03* | -0.24 | 0.48 | -0.32 | 0.26 |
|  | Loaded | -0.34 | 0.01* | -0.42 | 0.00* | -0.34 | 0.01* | -0.34 | 0.01* |
| November | Empty | -0.55 | 0.00* | -0.30 | 0.06 | -0.47 | 0.00* | -0.59 | 0.00* |
|  | Half loaded | -0.09 | 0.17 | -0.32 | 0.02* | -0.36 | 0.01* | -0.31 | 0.01* |
|  | Loaded | -0.16 | 0.24 | -0.51 | 0.03* | -0.49 | 0.04* | -0.52 | 0.02* |

\* Statistical significance at $p < 0.05$

There was a correlation between vegetation and photosynthetic indices calculated from the reflected spectra from tree canopies and the sugars content in the fruit (Figure 5). The four sugars in avocado fruit relate to each other positively along the season. There is a positive relationship between fructose, glucose and perseitol and to lesser extent between fructose, sucrose and D-mannoheptulose (Figure 5). As for the vegetation indices, NDVI is correlated with MTVI2 as expected, because both are related to the canopy structure. The photosynthetic vegetation indices SIF760 and SIF687 are also correlated between themselves because both relate to PhotoSystem II (PSII)’s LHC (Franck et al., 2002). SIF760 correlates better to the four sugars than SIF687.

**Figure 5.**
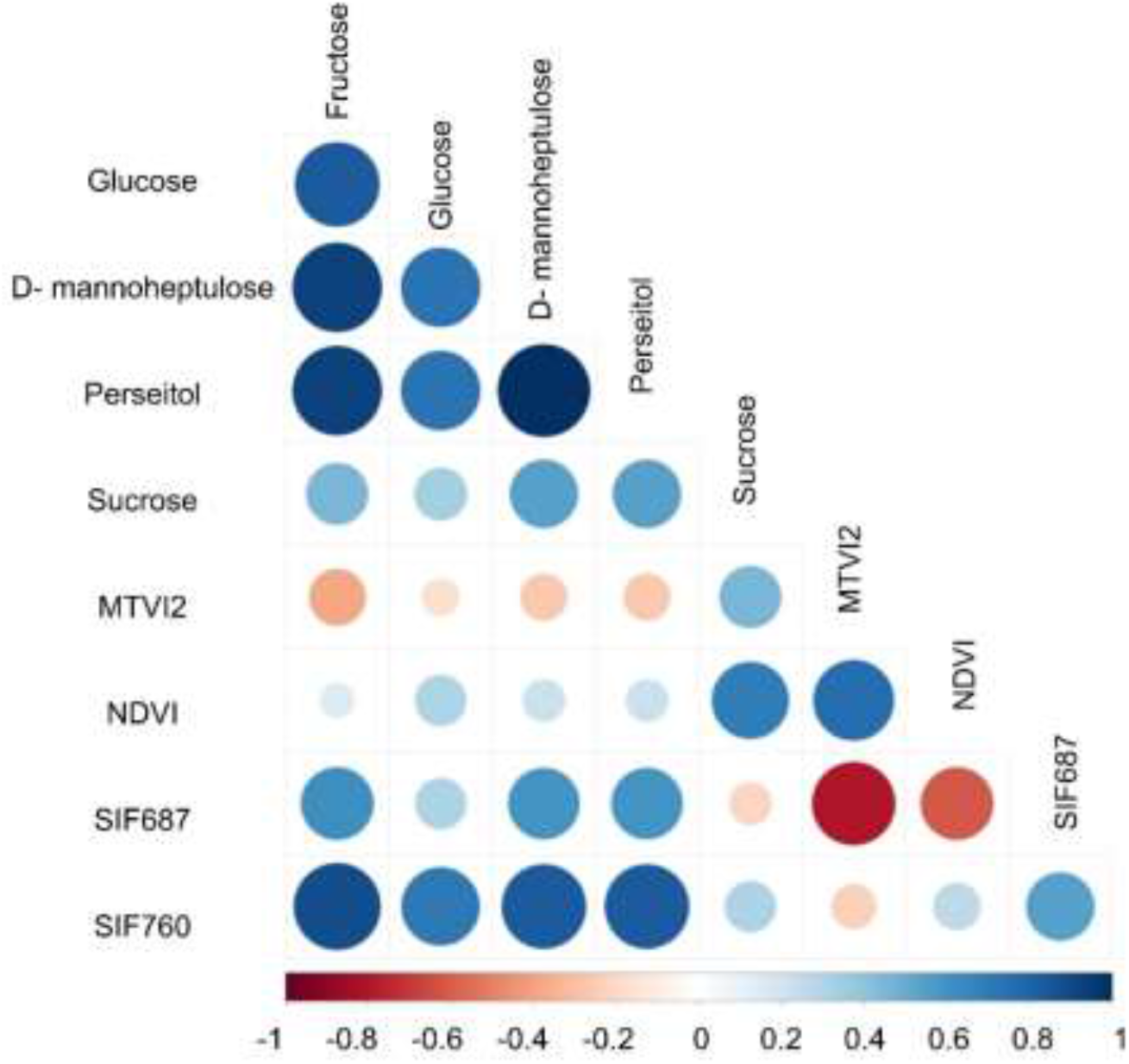
Correlation plot of photosynthetic and vegetation indices with accumulated sugars in avocado fruits. Higher correlation values between each two variables are presented as larger and darker symbols. Blue and red colors represent positive and negative correlation direction, respectively. N=50, omitting only the autocorrelated group samples.

The relationships between selected sugars and the photosynthesis indices are presented in (Figure 6). The sugars concentration was transformed to natural logarithm (Figure 4). The chlorophyll fluorescence indices demonstrate a non-linear relationship with the contents of perseitol and fructose, and the power dependent relationship revealed high correlation (Figure 6). The relationships of SIF687 and persitol content (Figure (6b)) obtains a similar behavior to that of SIF760 and fructose (Figure (6a)), however the strength of the nonlinear power relationship is not as high as for fructose and SIF760.

**Figure 6.**
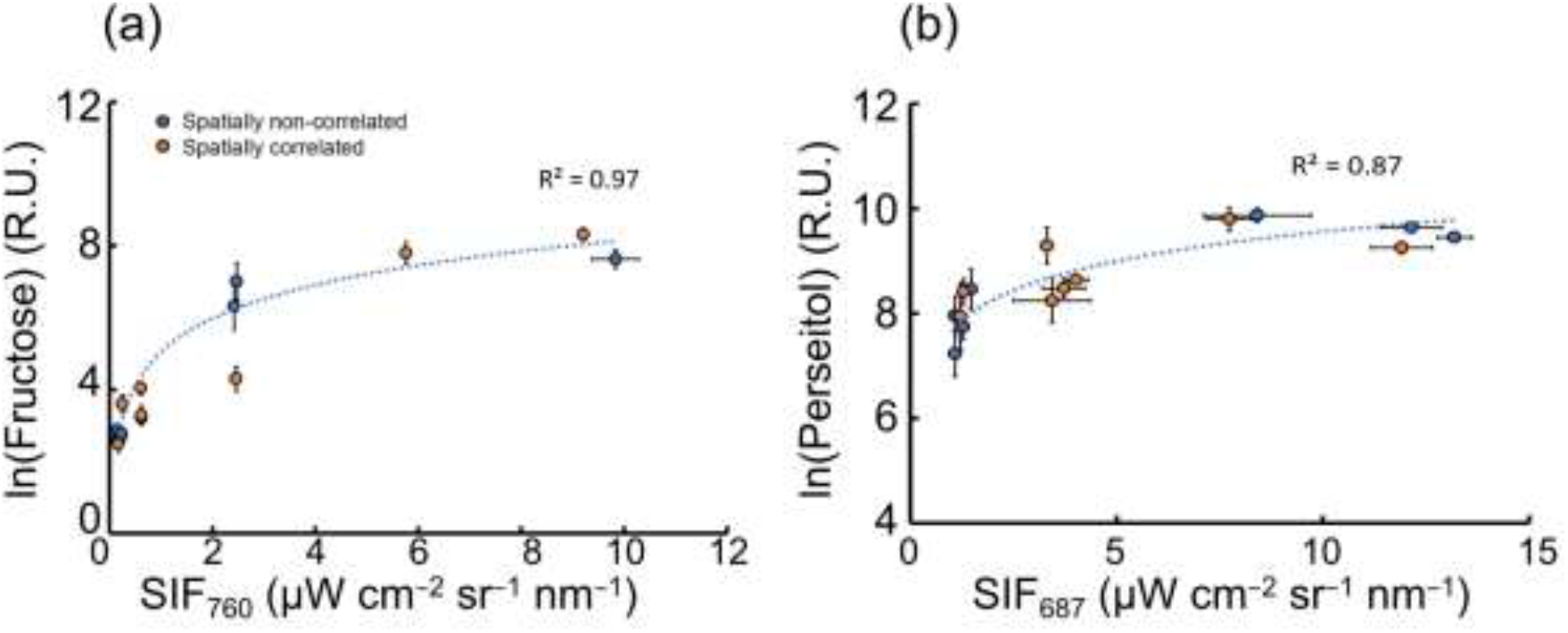
The relations between remote sensing indices of SIF and sugars content in ‘Hass’ avocado fruits. Panels (a) and (b) represent non-linear correlations between SIF and the natural logarithm of Fructose and Perseitol, respectively. N>3. Spatially non-correlated and correlated groups according to the Moran’s test are colored in blue and orange, respectively. Error bars represent standard error of the mean. Dotted curve colored in blue represents the overall correlation trend, including only the spatially non-correlated points.

More spatially autocorrelated groups were found towards the end of the season where sugar levels are depleted (the four orange points next to the y-axis in Figure (6a)). Autocorrelated groups are disperse evenly along the gradient of the perseitol (Figure (6b)) and not as seen for the fructose. All the measurement groups (crop load x sampling dates) are presented in Figure 6, where the power regression was performed only on groups that were not autocorrelated, in order not to violate the relation between independent samples when performing statistical regression (blue points curves in Figure 6 and Table 2).

## Discussion

Remote sensing of photosynthetic activity is focused mainly on crops and plants with homogenous canopy appearance: broadleaves, tropical and boreal forests, grasslands and wetlands, and field crops (Baldocchi et al. 2001). The literature obtains limited number of cases which relate between inhomogeneous canopy photosynthesis signals such as occurs on trees to actual leaf photosynthesis parameters (Wyber et al. 2018). In this study we used an interpolation technique that allows the calculation of SIF and vegetation indices for each tree in an avocado orchard. The tree-scale SIF values were correlated with sugar contents in avocado fruit pericarp. These correlations were apparent in spite the fact that in the case of *Persea Americana Mill.* the fruits are located within the canopy; thus, they are not visible to the sensor’s eye. With the level of sugars within the fruit that deplete over time until harvest (Delisiosa 1992) it is tempting to correlate between photosynthetic signals retrieved with remote sensing techniques and the dynamics of the sugar content within the fruit. Similarly, Shezi et al. 2019 shows that along the season chlorophyll fluorescence parameters measured at the outer side of the canopy of avocado trees at leaf level were connected to fruit maturity. This study relates between tree canopy photosynthesis activity and actual photosynthetic product, i.e. the result of biomass production. The dramatic reduction in sugar contents (Figure 4) is supported by other studies in avocado (Xuan Liu et al. 1999) (Kilaru et al. 2015), showing that within four months after June’s drop, avocado fruit sugars degrade and lose almost 50% of the total soluble sugar content. The dramatic reduction in sugar content in the fruit pericarp is related to maturation processes (Xuan Liu et al. 1999), and canopy fluorescence emission in avocado was related to fruit maturation (Shezi et al., 2019). Altogether, it seems that the dynamics of canopy photosynthesis indices along the season might be used to predict maturation time.

SIF687 was negatively correlated with MTVI2 (Figure 5), corroborating the fact that an increase in canopy cover reduces the probability of red photon to escape and not to be re-absorbed by the canopy (Van der Tol et al. 2009). Therefore, with increase in canopy there is a higher probability that emitted fluorescence at 687 nm will be absorbed back to the leaves, therefore less “red photons” will reach the sensor. SIF760 shows a very high correlation with all the sugars except sucrose (Figure 5). SIF760 is considered a much more stable signal that correlates with gross primary production better than SIF687 (Porcar-Castell, 2014). (Goulas et al. 2017) show that SIF760 was more correlated than SIF687 to biomass production rate in wheat, corroborating our findings that SIF760 correlates better to sugars synthesis. Part of this phenomenon lies in the fact that SIF687 is emitted solely from photosystem II (Franck, Juneau, and Popovic 2002), while SIF760 obtain a component of fluorescence emission from PSI as well. For this reason, SIF687 is a less stable signal due to photoprotective mechanisms which may attenuate the fluorescence signal emitted from PSII (Muller, Li, and Niyogi 2001).

SIF signal in this study was extracted according to the calculation of (Alonso et al. 2008), without correcting for reflectance entanglement from the reflected radiant flux. It may generate a limitation in various conditions (Meroni et al. 2009; Hikosaka and Tsujimoto 2021). However, Plascyk et al. (1975), which formulated the original SIF calculation, state that high quantum efficiency of the target will increase the stability of the signal, which is the situation in terrestrial plants in particular, and in photosynthetic organisms in general (Björkman and Demmig 1987). In addition, studies which use complex methodologies to correct for the reflected light may affect the SIF calculations, and do not necessarily show an improved extraction when compared to studies that did not used this methodology (Hikosaka and Tsujimoto 2021; Xinjie Liu et al. 2019; P.J. Zarco-Tejada, González-Dugo, and Fereres 2016).

Additional limitation is with the NDVI calculation which came out low for trees that were found on the borders of the plot where the pass from soil to canopy. This resulted in false mixed soil and canopy NDVI values. It relates to the size of the area covered by each spectroradiometer reading (6.5 m in diameter, whereas the tree crown was approximated at 4 m). In the current study the trees sampled for the measurement of sugar contents in the pericarp were not close to the orchard border. We suggest that if commercial use of the techniques employed here will be considered, the first and the last two trees in each row will be excluded from the indices mapping.

### Future perspectives

In order to validate the suggested methodology while interpreting signals from a UAV-SIF system, the authors suggest to perform a comparative study between simple and complex calculations of the SIF signal (with and without consideration of the reflected light entangled in the acquisition, respectively) (Meroni et al. 2009). This will establish rules to what extent each correction is necessary. Finally, this study suggests two future experiments in order to scale up measurements to a global scale: 1. Enabling a comparison between reflectance-based vegetation indices to the SIF signal at a high spatial and temporal resolution; 2. Comparing analysis in high spatial resolution to analysis of the same areas acquired by satellite with low resolution SIF and vegetation indices.

## Summary

In this study we retrieved SIF of an avocado orchard using a very high spectral resolution spectroradiometer mounted on a UAV, in order to track changes in avocado fruits sugar content during the main fruit development period. A computer algorithm was used to match between time stamps of the sensor and the position of the drone in time and space, and kriging interpolation provided a continuous approximation of the spectral vegetation indices allowing the calculation of the spectral indices for each tree in the orchard. The canopy SIF indices showed non-linear correlation with the sugar content in the avocado fruits, which are located within the canopy (not exposed to the sensor). It may serve as a basis for the development of vegetation indices originate from SIF to be used for precision agriculture decisions.

